# A reference genome for the Chinese Lizardtail Herb (*Saururus chinensis*)

**DOI:** 10.1101/2024.02.12.579984

**Authors:** Chengyi Tang

## Abstract

Several months earlier, other researchers had achieved the inaugural publication of the Chinese Lizardtail Herb (*Saururus chinensis*) genome dataset. However, the quality of that genome dataset is not deeply satisfactory, especially in terms of genome continuity (Contig N50 length ≈ 1.429 Mb) and gene-set completeness (BUSCO evaluation ≈ 91.32%). In this study, we present an improved chromosome-level genome of *S. chinensis*, characterized by heightened genome continuity (Contig N50 length ≈ 4.180 Mb) and a more complete gene-set (BUSCO evaluation ≈ 95.91%). Our investigation reveal that the extant *S*. chinensis genome preserves abundant vestiges of a paleo-tetraploidization event that are discernible both at the macroscopic chromosome level and within microscopic gene families, such as the PEL (pseudo-etiolation in light) family. Moreover, we elucidate that this paleo-tetraploidization event is associated with an expansion of the PEL family, potentially initiating a process conducive to its neofunctionalization and/or subfunctionalization.

## Introduction

The Chinese lizardtail herb (*Saururus chinensis*) (Fig. 1A), also commonly known as “Sanbaicao” within the scope of traditional Chinese medicine, is not only a well-known traditional herb used as a treatment for conditions such as edema, asthma, jaundice, gonorrhea and various other ailments, but it is also a core species within the family Sauraceae, in the order Piperales (Liu et al., 2020). Currently, the Piperales is divided into three families, namely Aristolochiaceae, Piperaceae, and Saururaceae (The Angiosperm Phylogeny Group, 2016). It is interesting to note that the two perianth-less (lacking petals and/or sepals) families, Piperaceae and Saururaceae, exhibit marked dissimilarity when juxtaposed with the perianth-bearing family Aristolochiaceae (Jaramillo et al., 2004; Remizowa et al., 2005). Due to its perianth-less floral composition and easy artificial propagation, *S. chinensis* has primarily found utility in genetic investigations to understand the origin of primitive flowering plants (Zhao et al., 2013; Zhao et al., 2021; Xue et al., 2023). More interestingly, *S. chinensis* owns prominent white bracts situated beneath its inflorescences, assuming a pivotal role in insect pollination (Song et al., 2018). Xue et al. (2023) have discerned that a specific PEL (pseudo-etiolation in light) gene from *S. chinensis*, namely ScPEL gene, inhibits chlorophyll biosynthesis and thus contribute to bract whitening. However, the understanding of the PEL family within the *S. chinensis* genome, as well as its evolutionary process, remains poorly elucidated.

**Fig. 1.**
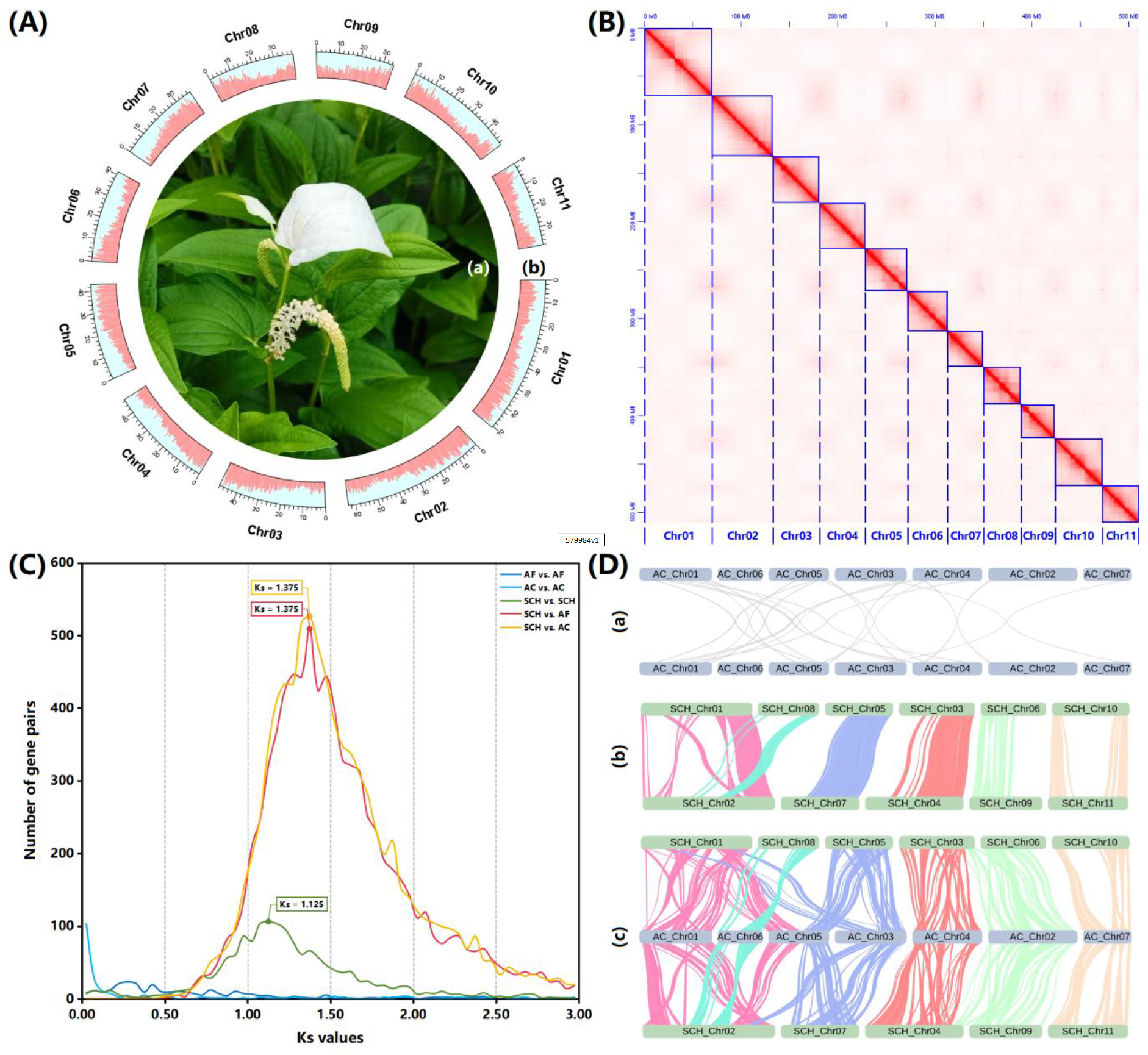
*S. chinensis* genome and its evolutionary process. (A) *S. chinensis* (a) and its genic distribution in chromosomes (b); (B) Chromosome interaction signals in *S. chinensis* (1n = 11); (C) Ks distribution curves (AF: *A. fimbriata*; AC: *A. contorta*; SCH: *S. chinensis*); (D) Chromosomal collinearity within *A. contorta* (a), within *S. chinensis* (b) and between *A. contorta* with *S. chinensis* (c).

To date, four genomes have been systematically reported within the aforementioned three families in the Piperales, namely *Aristolochia fimbriata* (Qin et al., 2021) and *A. contorta* (Cui et al., 2022) in Aristolochiaceae, *Piper nigrum* (Hu et al., 2019) in Piperaceae, and *S. chinensis* (Xue et al., 2023) in Saururaceae. It is important to note that *P. nigrum* and *S. chinensis* exhibit discernible low levels of gene-set completeness with respect to *A. fimbriata* and *A. contorta. P. nigrum* falls below 85% in gene-set completeness and *S. chinensis* is marginally above 90%, while *A. fimbriata* and *A. contorta* surpass 95% observably (Supplementary Table S1). Additionally, *P. nigrum* and *S. chinensis* exhibit significantly lower levels of genome continuity than *A. fimbriata* and *A. contorta*. The Contig N50 lengths (a statistical indicator of genome continuity) of *P. nigrum* and *S. chinensis* remain notably below 1.5 Mb, while those of *A. fimbriata* and *A. contorta* exceed 2.0 Mb, with that of *A. fimbriata* in particular exceeding 5.0 Mb (Supplementary Table S1). Thus, the state of the published genomes within the perianth-less families (Piperaceae and Saururaceae) highlights an inadequacy in meeting current scholarly expectations.

In the present study, we use short Illumina and long ONT genome sequencing plus Hi-C (high-through chromosome conformation capture) sequencing to obtain an updated chromosome-level genome of *S. chinensis* with enhanced genome continuity and a more complete gene-set. We believe that this reliable genome dataset could provide an effective resource for comparative genomics and molecular evolution research.

## Results and Discussion

### *S. chinensis* genome is a potential paleo-tetraploid

A comprehensive dataset for the *S. chinensis* genome was acquired, comprising ∼22.95 Gb of Illumina reads, ∼66.56 Gb of ONT reads, and ∼40.50 Gb of Hi-C reads (Supplementary Table S2). The estimated genome size of *S. chinensis* stood at ∼528.084 Mb (Supplementary Table S3), a concordance with the result reported by Xue et al. (2023). The final assembled genome spanned 522.247 Mb, exhibited a Scaffold N50 length of 46.948 Mb and a Contig N50 length of 4.180 Mb (Supplementary Table S1), and revealed the presence of eleven chromosomes (Fig. 1B), in agreement with previous investigations (Okada, 1986; Xue et al., 2023).

Subsequent annotations indicated that the genome comprised ∼275.71 Mb of repetitive sequences (∼52.81% of total genome), consisting of ∼211.583 Mb of interspersed repeats (∼40.52%), ∼4.120 Mb of tandem repeats (∼0.79%), and ∼58.794 Mb of unknown repeats (∼11.26%) (Supplementary Table S4). In addition, the genome comprised a total of 32,124 protein-coding gene models (Fig. 1A and Supplementary Table S1). BUSCO evaluation (Manni et al. 2021) demonstrated that ∼95.91% of the complete BUSCOs were aligned with total gene models (Supplementary Table S1), indicating an acceptable level of gene-set completeness. In summary, we had obtained an enhanced chromosome-level genome for *S. chinensis*, which was characterized by heightened genome continuity and a more complete gene set (Supplementary Table S1), surpassing the previous version (Xue et al., 2023).

The Ks distribution curve delineated that Ks_(SCH_vs_AF)_ (1.375) = Ks_(SCH_vs_AC)_ (1.375) > Ks_(SCH_vs_SCH)_ (1.125) (Fig. 1C), suggesting that *S. chinensis* underwent an ancient polyploidization event subsequent to its divergence from two *Aristolochia* species. This proposition was supported by the findings reported by Xue et al. (2023). Moreover, the present *S. chinensis* genome was obviously divided into two groups: Group A, consisting of chromosomes 1, 8, 5, 3, 6, and 10, and Group B, consisting of chromosomes 2, 7, 4, 9, and 11 (Fig. 1D), elucidating that this event manifested itself as a tetraploidization event. In addition, previous studies had highlighted that two *Aristolochia* species were typically diploid and had not undergone a polyploidization event after the ancestral ε event (Jiao et al., 2011; Qin et al., 2021; Cui et al., 2022). Our results also supported this proposition, which was evident at both the Ks distribution and genome collinearity levels (Fig. 1C-1D). Consequently, it became apparent that *S. chinensis*, after the tetraploidization event, still retained abundant vestiges of genome duplication from that epoch, and that it had not been thoroughly re-diploidized even to the present day. In other words, at the macroscopic chromosome level, the *S. chinensis* genome could be referred to as a “paleo-tetraploid”. More interestingly, several genome fusions occurred following the paleo-tetraploidization event within the *S. chinensis* genome. A prominent illustration was the chromosome-level fusion of collinear blocks associated with the current Chromosome No. 8 into the central region of Chromosome No. 2 (Fig. 1D).

### Paleo-tetraploidization promotes PEL family expansion in *S. chinensis*

We discerned a singular complete domain, namely “PF09713|A_thal_3526” (https://www.ebi.ac.uk/interpro/entry/pfam/PF09713/), within the ScPEL protein, indicating its distinctive structural feature (Supplementary Table S5). Using domain plus sequence similarity searches, we found that the ScPEL gene is equivalent to the SCH04C1170-RA gene in our *S. chinensis* genome (Supplementary Table S6), and further identified a total of six PEL genes in the *S. chinensis* genome, four in the *A. fimbriata* genome, and three in the *A. contorta* genome (Fig. 2A and Supplementary Table S5-S6). A phylogenetic analysis delineated the PEL family into four distinct clades (Fig. 2A). There also were discernible differences in the length of amino acid sequences corresponding to the four clades, with Clade III exhibiting the highest length, followed by Clade IV, Clade II, and Clade I (Fig. 2A). Interestingly, a quantitative expansion of the PEL genes was observed in *S. chinensis* compared to two *Aristolochia* species. This genic expansion was attributed to the paleo-tetraploidization event that occurred within the *S. chinensis* genome (Fig. 1C-1D), resulting in gene duplication in Clades I and III (Fig. 2A), and also revealing the enduring imprint of the paleo-tetraploidization event on the genic landscape at the microscopic scale (Fig. 2B). This expansion augmented genic resources, potentially contributing to the neofunctionalization and/or subfunctionalization of the PEL family within the *S. chinensis* genome, and thus to the origin of the distinctive white bracts.

**Fig. 2.**
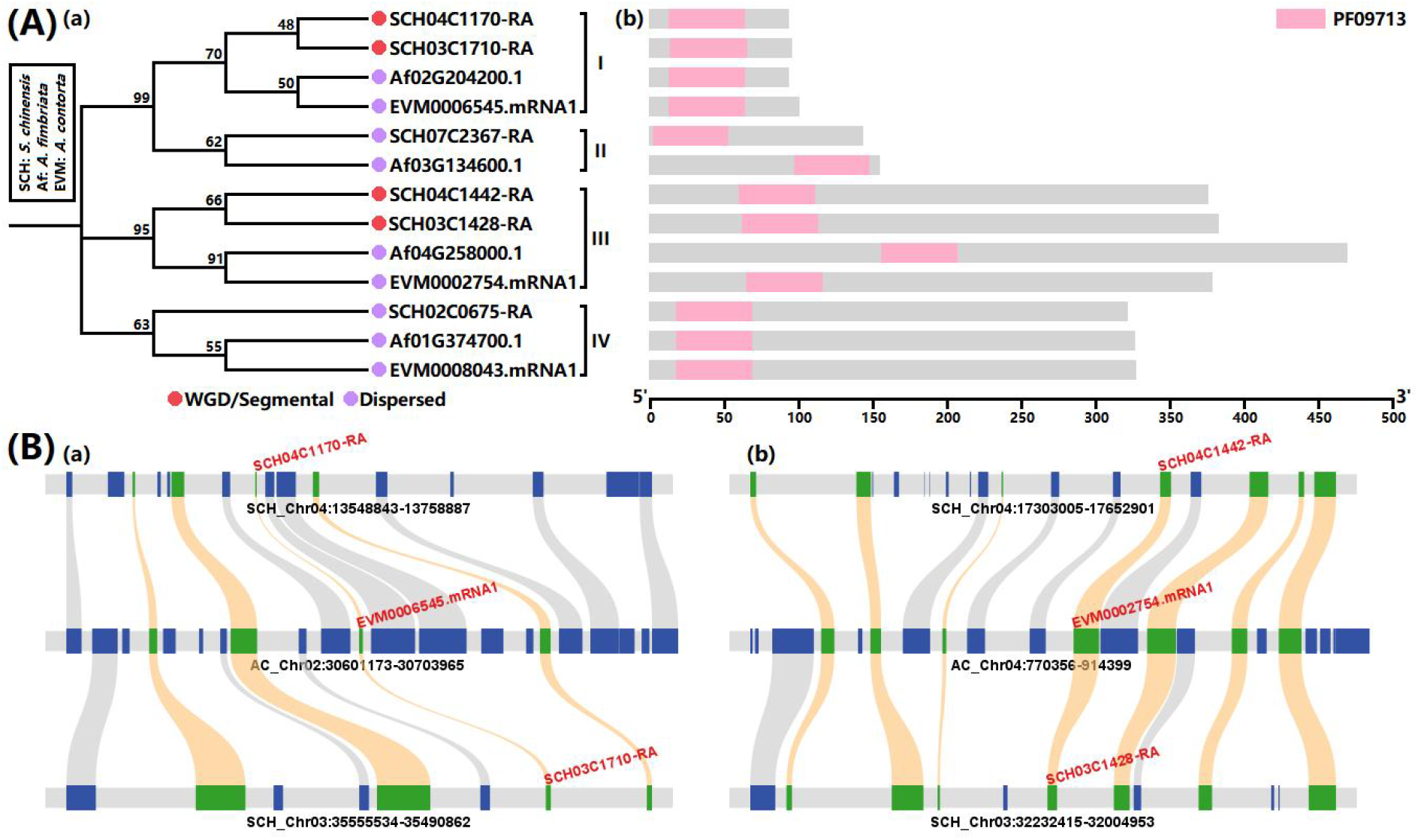
PEL family and its evolutionary process. (A) Phylogenetic structure (a) and sequence characteristics (b) of PEL family; (B) Microscopic collinearity of PEL family expansion (a. Clade I and b. Clade III) between *A. contorta* with *S. chinensis*.

## Materials and Methods

### Plant materials

Initially, some Chinese lizardtail herb seedlings were procured from Shenzhen Yuanzhihui Company through the 1688 online trading platform (https://shop3i77d84502842.1688.com). Subsequently, these seedlings underwent cultivation within a greenhouse environment (Fig. 1A), maintaining a temperature of ∼25°C and a photoperiod of 14 hours of light contrasted with 10 hours of darkness. After approximately 60 days, fresh leaves obtained from a vigorous individual were collected for genome and Hi-C sequencing. Simultaneously, fresh leaves and stems from the same individual were collected for transcriptome sequencing.

### Genome, Hi-C and transcriptome sequencing

Total DNAs were extracted using the Magnetic Plant Genomic DNA Kit (Cat. no: 4992407; Tiangen, China). A paired-end library with an insert size of 350 kb was constructed using the TIANSeq Fast DNA Library Kit (Cat. no: 4992261; Tiangen, China) and then sequenced using an Illumina NovaSeq6000 sequenator (Illumina, USA). An amplification-free whole genome sequencing library was constructed using the Ligation Sequencing Kit (No: SQK-LSK110; ONT, UK) and then sequenced by a PromethION sequenator (ONT, UK). A Hi-C library was constructed following a recognized Hi-C protocol (restriction enzyme: HindIII) described in earlier studies (Grob et al., 2014; Rao et al., 2014; Hu et al., 2019; Tang et al., 2020; Qin et al., 2021; Cui et al., 2022; Xue et al., 2023) and then sequenced by an Illumina NovaSeq6000 sequenator (Illumina, USA). Total RNAs were extracted using an RNAprep Pure Plant Kit (Cat. no: 4992237; Tiangen, China). A cDNA library was constructed using the TIANSeq Fast RNA Library Prep Kit (Cat. no: 4992376; Tiangen, China) and then sequenced using an Illumina NovaSeq6000 sequenator (Illumina, USA).

### Data processing

Fastp v0.23.2 (Chen et al., 2018) filtered Illumina raw data to remove adapters, low-quality reads, and poly-N reads. NanoFilt v2.8.0 (De Coster et al., 2018) filtered ONT raw data to remove too-short reads (i.e., length < 2 kb) and low-quality reads (i.e., RQ < 7.0).

### Genome size estimation

The Illumina clean data (Supplementary Table S2) was applied for genome size estimation. K-mers were counted and exported to a histogram file using Jellyfish v2.3.0 (key parameters: jellyfish count -m 17, 19 or 21; jellyfish histo -h Max_count) (Marcais and Kingsford, 2011).

Preliminary genome sizes were calculated using GenomeScope 2.0 (Ranallo-Benavidez et al., 2020) or GCE v1.0.2 (Liu et al., 2013), and the final genome size was averaged over the preliminary genome sizes.

### Genome assembly

The ONT clean data (Supplementary Table S2) was used for genome assembly. First, NextDenovo v2.5.0 (key parameters: seed_depth = 999, nextgraph_options = -a 1 -u 1 -G) (Hu et al., 2023) was executed for contig-level assembly. Over-redundant contigs were removed via purge_dups v1.2.6 (key parameters: -a 50) (Guan et al., 2020). NextPolish v1.4.1 (Hu et al., 2020) was then used for genome polishing based on ONT and Illumina clean data (key parameters: task = 551212). Subsequently, the Hi-C clean data (Supplementary Table S2) was mapped to this polished contig-level assembly using Juicer v1.6 (key parameters: -s HindIII) (Durand et al., 2016a) and ordered to a chromosome-level assembly via 3D-DNA (Dudchenko et al., 2017). Finally, Juicebox v1.11.08 (Durand et al., 2016b) was used to manually curate this chromosome-level assembly.

### Genome annotation

Repetitive sequences were annotated via RepeatMasker v4.1.4 (http://www.repeatmasker.org), based on a combined database including Dfam v3.7 (Storer et al., 2021) plus a *de novo* custom library of *S. chinensis* constructed via RepeatModeler v2.0.4 (key parameter: -LTRStruct) (Flynn et al., 2020). Subsequently, protein-coding genes were annotated as the following process: (1) Repetitive sequences were masked first; (2) AUGUSTUS v3.5.0 (Stanke et al., 2008) and GeneMark-EP+ v4.71 (Bruna et al., 2020) were used for *ab-initio* predictions; (3) Exonerate v2.4.0 (Slater and Birney, 2005) was applied to homological predictions based on published genomes of two related species, i.e., *A. fimbriata* (Qin et al., 2021) and *A. contorta* (Cui et al., 2022); (4) PASA v2.5.2 (Haas et al., 2021) was used to identify transcripts based on the transcriptome data (Supplementary Table S2); (5) The total results were integrated into a joint gene-set using Maker v3.01.03 (key parameters: softmask = 1; min_protein = 49) (https://www.yandell-lab.org/software/maker.html). Moreover, BUSCO v5.2.2 (Manni et al. 2021) was used to evaluate gene-set completeness (key parameters: -l embryophyta_odb10 -m proteins).

### Comparative genome analysis

Gene sets were employed for sequence similarity search via BLASTP 2.13.0+ (key parameter: -evalue 1e-5 -max_target_seqs 5) (Camacho et al., 2009). Subsequently, the obtained results were further analyzed using MCScanX (Wang et al., 2012) to identify collinearity within monophyletic or crossed species. In addition, all gene pairs within the identified collinear blocks were aligned individually via MUSCLE v3.8.31 (Edgar, 2004), predicated on their amino acid sequences, and all alignments were then transformed back to nucleotide sequences. The computation of Ks values for individual gene pairs was performed via KaKs_Calculator 3.0 (Zhang, 2022), based on these nucleotide alignments. The peaks in the Ks distribution curves of monophyletic species were indicative of polyploidization events, while the peaks in the Ks distribution curves of crossed species were indicative of divergence events.

### Gene family analysis

Initially, the hmmscan module (key parameter: -E 1e-05) within HMMER 3.3.2 (Potter et al., 2018) was employed for domain identification in the ScPEL protein. Subsequently, utilizing the PF09713 domain as a query, domain similarity search was performed via the hmmsearch module (key parameter: -E 1e-05) within HMMER 3.3.2 (Potter et al., 2018). Concurrently, utilizing the ScPEL protein as a query, sequence similarity search was performed via BLASTP 2.12.0+ (key parameters: -evalue 1e-5 -max_target_seqs 500) (Camacho et al., 2009). The redundant PEL members were removed from the final output set. Following this, the identified PEL proteins were aligned via MUSCLE v3.8.31 (Edgar, 2004). The initial alignment was trimmed via trimAl v1.4.1 (key parameter: -gt 0.50) (Capella-Gutierrez et al., 2009). The trimmed alignment was used as the basis for constructing a phylogenetic tree via IQ-TREE v2.2.2 (best-fit model: JTT+I+G4; key parameters: --seqtype AA -m MFP --alrt 1000 -B 1000) (Minh et al., 2020) according to the ML (maximum likelihood) method. The delineation of the PEL family was based on the hierarchical structure of the phylogenetic tree. Additionally, the ‘duplicate_gene_classifier’ module within MCScanX (Wang et al., 2012) was utilized to determine the duplication types for each PEL gene.

## Acknowledgments

Thanks for financial supports from the National Natural Science Foundation of China (Grant Numbers: 32000400) and the Natural Science Foundation of Jiangsu Province (Grant Numbers: BK20180332).

## Data Availability

All sequencing data from this study have been stored in NCBI BioProject (Accession No.: PRJNA1066966) and their detailed SRA IDs are listed in Supplementary Table S2. *Saururus chinensis* genome and its gene set have been made publicly accessible on figshare (Download Link: https://doi.org/10.6084/m9.figshare.25035707.v1).

