## Supplementary Tables S1-S6 for "A reference genome for the Chinese Lizardtail Herb (*Saururus chinensis*)"

**Supplementary Table S1. Summary statistics of published genomes within Piperales (Up to 2024.01.15).**

| Statistical types | *A. fimbriata* | | *A. contorta* | | *P. nigrum* | | *S. chinensis* | | *S. chinensis* | |
|---|---|---|---|---|---|---|---|---|---|---|
| | Scaffolds | Contigs | Scaffolds | Contigs | Scaffolds | Contigs | Scaffolds | Contigs | Scaffolds | Contigs |
| Total sequences | 232 | 1,065 | 84 | 224 | 45 | 1,300 | 38 | 842 | 72 | 323 |
| Total lengths (bp) | 257,701,785 | 249,415,861 | 210,540,771 | 210,526,771 | 761,218,309 | 756,720,420 | 539,018,180 | 538,937,780 | 522,246,995 | 522,221,895 |
| Gap lengths (bp) | 8,285,924 | 0 | 14,000 | 0 | 4,497,889 | 0 | 80,400 | 0 | 25,100 | 0 |
| Maximum lengths (bp) | 45,753,714 | 10,542,392 | 38,743,438 | 12,320,100 | 48,451,882 | 5,646,249 | 70,783,412 | 5,467,227 | 70,549,925 | 20,829,987 |
| N50 lengths (bp) | 33,250,414 | **5,157,218** | 30,347,173 | **2,315,928** | 29,812,524 | **1,092,847** | 47,843,222 | **1,428,727** | 46,947,598 | **4,179,527** |
| N90 lengths (bp) | 22,664,973 | 327,827 | 19,489,044 | 388,540 | 22,167,516 | 298,494 | 39,887,764 | 314,759 | 37,456,933 | 1,170,872 |
| Total gene models | 21,751 | | 18,315 | | 63,466 | | 36,140 | | 32,124 | |
| BUSCO evaluation for gene-set (embryophyta_odb10, 1614) | **C:96.72%** [S:94.73%, D:1.99%], F:1.24%, M:2.04% | | **C:96.22%** [S:90.58%, D:5.64%], F:1.74%, M:2.04% | | **C:82.28%** [S:58.24%, D:24.04%], F:7.00%, M:10.72% | | **C:91.32%** [S:87.17%, D:4.15%], F:5.27%, M:3.41% | | **C:95.91%** [S:92.87%, D:3.04%], F:1.12%, M:2.97% | |
| Sources | Qin et al. (2021) | | Cui et al. (2022) | | Hu et al. (2019) | | Xue et al. (2023) | | This study | |

**Supplementary Table S2. Summary statistics of sequencing data from this study.**

| Sequencing types | Sequencing platforms | Samples | Clean data size (Gb) | NCBI SRA IDs |
|---|---|---|---|---|
| Genome | Illumina NovaSeq6000 | Fresh leaves | 22.95 | SRR27665351 |
| | ONT PromethION | Fresh leaves | 66.56 | SRR27686889 |
| Hi-C | Illumina NovaSeq6000 | Fresh leaves | 40.50 | SRR27673968 |
| Transcriptome | Illumina NovaSeq6000 | Fresh leaves and stems | 4.81 | SRR27669428 |

**Supplementary Table S3. Genome size estimation of *S. chinensis* in this study.**

| K values | Preliminary size (Mb) | | | Final average size (Mb) |
|---|---|---|---|---|
| | GenomeScope v2.0 | GCE v1.0.2 | Software average | |
| 17 | 522.223 | 502.110 | 512.167 | 528.084 |
| 19 | 523.091 | 553.490 | 538.291 | |
| 21 | 520.604 | 546.985 | 533.795 | |

**Supplementary Table S4. Summary statistics of repetitive sequences within *S. chinensis* genome from this study.**

| Statistical types | | | | Size (bp) | Percent (%) |
|---|---|---|---|---|---|
| Interspersed repeats | RNA transposons | LTRs | *Copia* | 38,124,324 | 7.3004 |
| | | | *Gypsy* | 102,417,503 | 19.6119 |
| | | | Unknown | 45,268,519 | 8.6684 |
| | | | Total | 185,810,346 | 35.5807 |
| | | LINEs | | 2,219,418 | 0.4250 |
| | | SINEs | | 6,257 | 0.0012 |
| | | Total | | 188,036,021 | 36.0069 |
| | DNA transposons | hAT | | 3,475,301 | 0.6655 |
| | | *Tc1/mariner*-like | | 27,283 | 0.0052 |
| | | *Harbinger*-like | | 971,885 | 0.1861 |
| | | Others and Unknown | | 18,346,976 | 3.5133 |
| | | Total | | 22,821,445 | 4.3701 |
| | Unknown | | | 725,044 | 0.1388 |
| | Total | | | 211,582,510 | 40.5158 |
| Tandem repeats | Simple repeats | | | 4,106,983 | 0.7864 |
| | Satellites | | | 13,082 | 0.0025 |
| | Total | | | 4,120,065 | 0.7889 |
| Others | | | | 1,274,426 | 0.2440 |
| Unknown | | | | 58,793,978 | 11.2584 |
| Total repetitive sequences (non-overlap) | | | | 275,770,664 | 52.8072 |

**Supplementary Table S5. Results of domain similarity searches for PF09713 domain.**

| Query ID | Subject IDs | e-values | Alignment information | | | |
|---|---|---|---|---|---|---|
| | | | Query start | Query end | Subject start | Subject end |
| PF09713 | **ScPEL** | **3.7E-27** | **1** | **52** | **12** | **63** |
| | SCH02C0675-RA | 6.5E-25 | 1 | 52 | 17 | 68 |
| | SCH03C1428-RA | 3.5E-25 | 1 | 52 | 61 | 112 |
| | SCH03C1710-RA | 5.2E-26 | 1 | 52 | 13 | 64 |
| | SCH04C1170-RA | 3.7E-27 | 1 | 52 | 12 | 63 |
| | SCH04C1442-RA | 6.8E-25 | 1 | 52 | 59 | 110 |
| | SCH07C2367-RA | 8.4E-23 | 1 | 51 | 2 | 52 |
| | Af01G374700.1 | 1.3E-25 | 1 | 52 | 17 | 68 |
| | Af02G204200.1 | 1.0E-26 | 1 | 52 | 12 | 63 |
| | Af03G134600.1 | 1.8E-21 | 1 | 51 | 96 | 146 |
| | Af04G258000.1 | 4.3E-25 | 1 | 52 | 154 | 205 |
| | EVM0002754.mRNA1 | 3.6E-25 | 1 | 52 | 64 | 115 |
| | EVM0006545.mRNA1 | 2.5E-27 | 1 | 52 | 12 | 63 |
| | EVM0008043.mRNA1 | 1.2E-25 | 1 | 52 | 17 | 68 |

**Supplementary Table S6. Results of sequence similarity searches for ScPEL protein.**

| Query ID | Subject IDs | e-values | Alignment information | | | | |
|---|---|---|---|---|---|---|---|
| | | | Identity (%) | Query start | Query end | Subject start | Subject end |
| ScPEL | SCH02C0675-RA | 7.4E-13 | 50.9 | 9 | 65 | 14 | 70 |
| | SCH03C1428-RA | 7.3E-13 | 45.2 | 9 | 70 | 58 | 119 |
| | SCH03C1710-RA | 2.7E-42 | 75.0 | 4 | 91 | 5 | 92 |
| | **SCH04C1170-RA** | **2.6E-65** | **100.0** | **1** | **92** | **1** | **92** |
| | SCH04C1442-RA | 4.6E-13 | 50.9 | 9 | 65 | 56 | 112 |
| | SCH07C2367-RA | 1.2E-22 | 69.2 | 11 | 62 | 1 | 52 |
| | Af01G374700.1 | 2.6E-14 | 52.6 | 9 | 65 | 14 | 70 |
| | Af02G204200.1 | 1.2E-42 | 73.6 | 1 | 91 | 1 | 91 |
| | Af03G134600.1 | 6.2E-20 | 58.9 | 7 | 62 | 91 | 146 |
| | Af04G258000.1 | 4.2E-13 | 46.8 | 9 | 70 | 151 | 212 |
| | EVM0002754.mRNA1 | 8.9E-13 | 50.9 | 9 | 65 | 61 | 117 |
| | EVM0006545.mRNA1 | 1.0E-42 | 72.5 | 1 | 91 | 1 | 91 |
| | EVM0008043.mRNA1 | 6.8E-14 | 52.6 | 9 | 65 | 14 | 70 |
